# Idiosyncratic purifying selection on metabolic enzymes in the long-term evolution experiment with *Escherichia coli*

**DOI:** 10.1101/2021.01.16.426962

**Authors:** Rohan Maddamsetti

## Abstract

Bacteria, Archaea, and Eukarya all share a common set of metabolic reactions. This implies that the function and topology of central metabolism has been evolving under purifying selection over deep time. Central metabolism may similarly evolve under purifying selection during longterm evolution experiments, although it is unclear how long such experiments would have to run (decades, centuries, millennia) before signs of purifying selection on metabolism appear. I hypothesized that central and superessential metabolic enzymes would show evidence of purifying selection in the long-term evolution experiment with *Escherichia coli* (LTEE). I also hypothesized that enzymes that specialize on single substrates would show stronger evidence of purifying selection in the LTEE than generalist enzymes that catalyze multiple reactions. I tested these hypotheses by analyzing metagenomic time series covering 62,750 generations of the LTEE. I find mixed support for these hypotheses, because the observed patterns of purifying selection are idiosyncratic and population-specific. To explain this finding, I propose the Jenga hypothesis, named after a children’s game in which blocks are removed from a tower until it falls. The Jenga hypothesis postulates that loss-of-function mutations degrade costly, redundant, and nonessential metabolic functions. Replicate populations can therefore follow idiosyncratic trajectories of lost redundancies, despite purifying selection on overall function. I tested the Jenga hypothesis by simulating the evolution of 1,000 minimal genomes under strong purifying selection. As predicted, the minimal genomes converge to different metabolic networks. Strikingly, the core genes common to all 1,000 minimal genomes show consistent signatures of purifying selection in the LTEE.

**Significance Statement:** Purifying selection conserves organismal function over evolutionary time. However, few studies have examined the role of purifying selection during adaptation to novel environments. I tested metabolic enzymes for purifying selection in an ongoing long-term evolution experiment with *Escherichia coli.* While some populations show signs of purifying selection, the overall pattern is inconsistent. To explain these findings, I propose the Jenga hypothesis, in which loss-of-function mutations first degrade costly, redundant, and nonessential metabolic functions, after which purifying selection begins to dominate. I then tested several predictions of the Jenga hypothesis using computational simulations. On balance, the simulations confirm that we should find evidence of purifying selection on the metabolic pathways that sustain growth in a novel environment.

## Introduction

Researchers have learned much about adaptive processes by conducting evolution experiments with large populations of microbes. By contrast, researchers have rarely, if ever, used experimental evolution to study purifying selection, because it may take hundreds or even thousands of years to see evolutionary stasis in action. My colleagues and I have recently started to address this research gap, by examining detailed time series of genetic change in Lenski’s long-term evolution experiment with *Escherichia coli* (LTEE). Our research addresses the following question: what processes are evolving under purifying selection, during adaptation to a novel laboratory environment in which abiotic conditions are held constant?

The LTEE has studied the evolution of 12 initially identical populations of *Escherichia coli* in carbon-limited minimal medium for more than 30 years and 60,000 generations of bacterial evolution (Lenski, et al. 1991; Lenski 2017). The LTEE populations are named by the presence of a neutral phenotypic marker: populations Ara+1 to Ara+6 grow on arabinose, while populations Ara-1 to Ara-6 cannot (Lenski, et al. 1991). The LTEE populations have diverged in their mutation rates and biases, as several LTEE populations have evolved elevated mutation rates due to defects in DNA repair (Sniegowski, et al. 1997; Tenaillon, et al. 2016; Good, et al. 2017; Maddamsetti and Grant 2020). These hypermutator populations tend to adapt faster than nonmutator populations that have retained the ancestral mutation rate (Wiser, et al. 2013), even though their more rapid genomic evolution largely reflects the accumulation of nearly neutral mutations through genetic hitchhiking (Couce, et al. 2017).

Because so many mutations are observed in the hypermutator populations, my colleagues and I hypothesized that sets of genes that are depleted of observed mutations may be evolving under purifying selection. To test this hypothesis, we developed a randomization test that we call STIMS (Simple Test to Infer Mode of Selection) to detect selection on pre-defined sets of genes, using metagenomic time series covering 60,000 generations of the LTEE (Good, et al. 2017). STIMS can detect both positive and purifying selection, and is able to control for both temporal variation in mutation rates over time as well as mutation rate variation over the chromosome (Maddamsetti and Grant 2020; Maddamsetti and Grant 2022). Using STIMS, we found evidence of purifying selection on aerobic-specific and anaerobic-specific genes (Grant, Maddamsetti, et al. 2021) and essential genes (Maddamsetti and Grant 2022) in the LTEE. In addition, we found evidence of purifying selection on protein interactome resilience (Maddamsetti 2021a), and purifying selection on genes encoding highly expressed and highly interacting proteins (Maddamsetti 2021b), using different methods.

Here, I use STIMS to test four sets of metabolic genes for purifying selection in the LTEE: (1) those that encode enzymes that catalyze the core reactions of *E. coli* central metabolism, as curated in the BiGG Models knowledge base (King, et al. 2015); (2) those encoding enzymes that catalyze “superessential” reactions that have been found to be essential in all tested bacterial metabolic network models (Barve, et al. 2012); (3) and (4) those that encode specialist and generalist enzymes, respectively, in the *E. coli* metabolic reaction network, as determined by their substrate and reaction specificity (Nam, et al. 2012).

I tested two specific hypotheses. First, I hypothesized that the BiGG core metabolic enzymes and the superessential metabolic enzymes would show evidence of purifying selection. Second, I hypothesized that specialist enzymes would show stronger purifying selection than generalist enzymes. My results show mixed support for these hypotheses. I find that patterns of purifying selection on metabolism, at least after 60,000 generations of experimental evolution, are largely idiosyncratic and population-specific. To explain this finding, I propose the Jenga hypothesis, in which mutation accumulation causes replicate populations to follow idiosyncratic trajectories of lost redundancies, despite purifying selection on network function. I tested several predictions of the Jenga hypothesis by simulating the evolution of 1,000 minimal genomes under strong purifying selection. Strikingly, the core genes common to all of the minimal genomes show consistent signatures of purifying selection in the LTEE. These signatures of purifying selection relate to the particular carbon sources that the LTEE populations are utilizing for growth.

## Results

### Overlap between pre-defined sets of metabolic genes

I first examined the overlap between the four gene sets of interest (Figure 1). If these sets are disjoint, then running STIMS on each would represent four independent statistical tests. At the other extreme, I would be conducting the same test four times.

**Figure 1.**
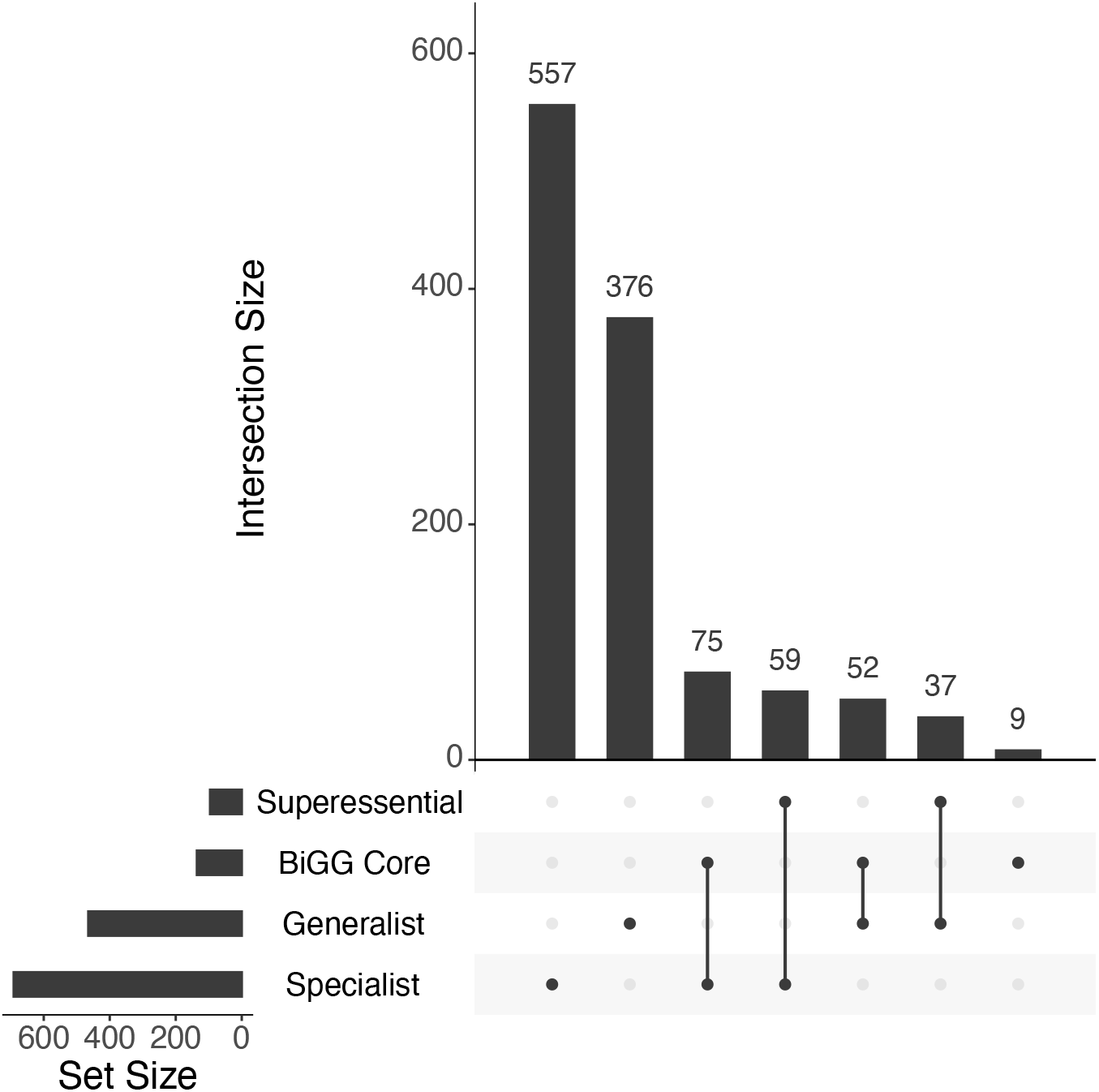
Intersections among the pre-defined sets of metabolic enzymes analyzed in this work. The intersections between four pre-defined sets: superessential enzymes, BiGG core metabolic enzymes, specialist enzymes, and generalist enzymes were examined (see text for further details about these gene sets). The cardinality of each of these sets is shown in the horizontal bar graph on the left. Every non-empty subset of the four sets is visualized as a set of dots connected by lines. The cardinality for each non-empty subset (set intersections) is shown in the vertical bar graph.

Specialist and generalist enzymes are mutually exclusive by definition. While I expected the BiGG core and superessential enzymes to overlap, due to their importance to *E. coli* metabolism, they are mutually exclusive (Figure 1). Out of 136 BiGG core enzymes, 75 are specialist enzymes, while 52 are generalist enzymes. The remaining 9 were not classified as either by Nam et al. (2012). Out of 96 superessential enzymes, 59 are specialist enzymes, and 37 are generalist enzymes. Therefore, two pairs of comparisons— BiGG core versus superessential, and specialist versus generalist— are statistically independent.

### Purifying selection on BiGG core metabolic enzymes

Out of the 6 hypermutator populations, 3 show purifying selection on the BiGG core metabolic genes: Ara-1 (STIMS bootstrap: *p* = 0.05), Ara+3 (STIMS bootstrap: *p* = 0.035), and Ara+6 (STIMS bootstrap: *p* = 0.003). In addition, Ara-4 trends toward purifying selection (STIMS bootstrap: *p* = 0.066). Therefore, the BiGG core metabolic genes show purifying selection in some—but not all—hypermutator LTEE populations (Figure 2A). No trend is seen in the nonmutator LTEE populations (Supplementary Figure 1A).

**Figure 2.**
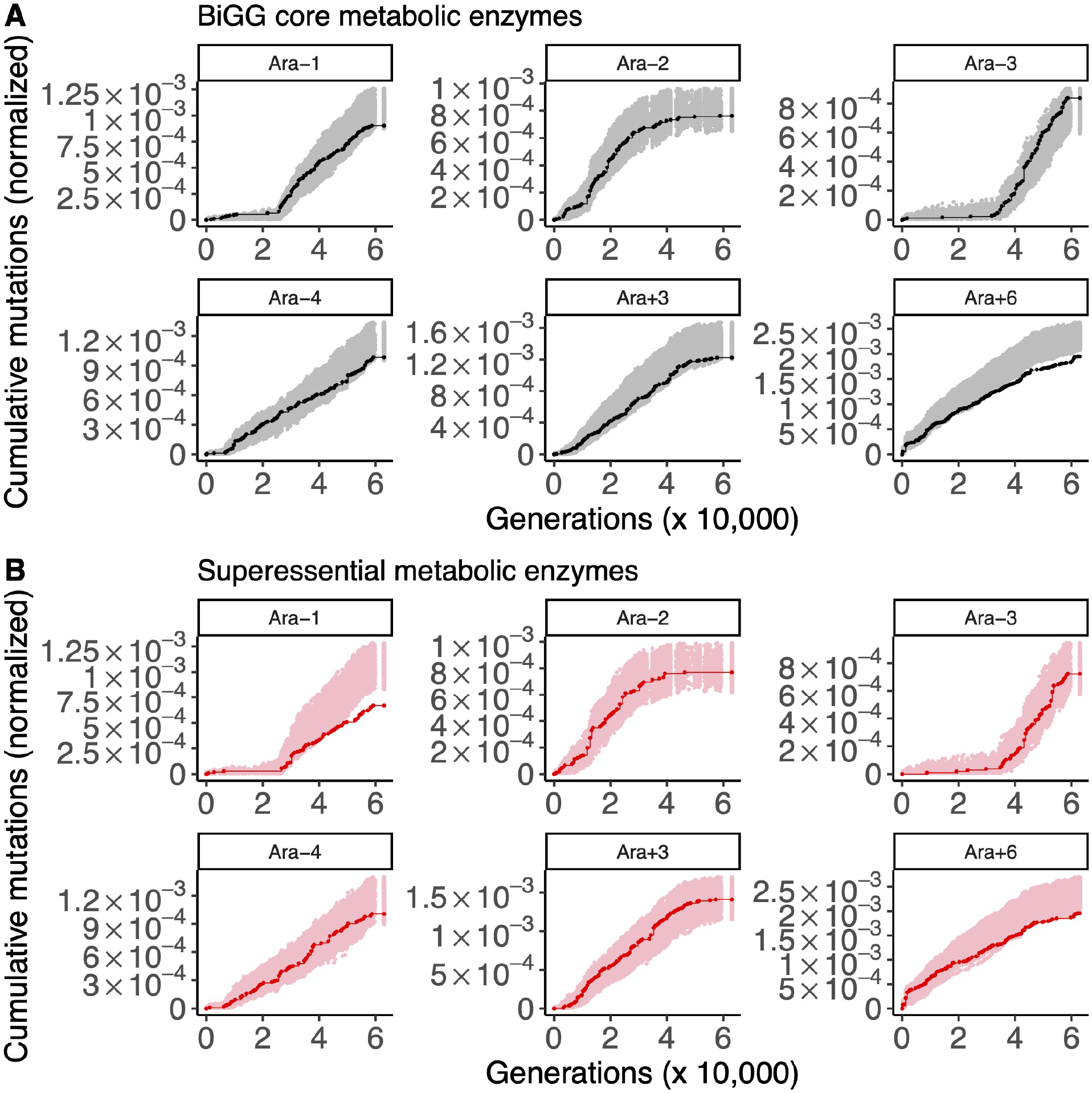
The results of running STIMS on BiGG core enzymes and superessential enzymes (hypermutator populations only). Each panel shows the cumulative number of mutations in the gene set of interest (solid line) in the six hypermutator LTEE populations. For comparison, random sets of genes (with the same cardinality as the gene set of interest) were sampled 1,000 times, and the cumulative number of mutations in those random gene sets, normalized by gene length, were calculated. The middle 95% of this null distribution is shown as shaded points. When a solid line falls below the shaded region, then the gene set of interest show a significant signal of purifying selection. A) The results of running STIMS on enzymes in the BiGG *E. coli* core metabolism model. B) The results of running STIMS on enzymes catalyzing superessential metabolic reactions.

### Idiosyncratic purifying selection on superessential metabolic genes

Only two populations show evidence of purifying selection on superessential metabolic genes: Ara-1 (STIMS bootstrap: *p* < 0.001) and Ara+6 (STIMS bootstrap: *p* = 0.01). The strength of purifying selection on superessential genes therefore seems to be population-specific (Figure 2B). No trend is seen in the nonmutator LTEE populations (Supplementary Figure 1B).

### Specialist enzymes tend to evolve under stronger purifying selection than generalist enzymes

Two hypermutator populations show evidence of purifying selection on specialist enzymes (Figure 3A): Ara–4 (STIMS bootstrap: *p* = 0.049) and Ara+6 (STIMS bootstrap: *p* < 0.001). Two nonmutator populations are significantly depleted of mutations in specialist enzymes (Supplementary Figure S2A): Ara+1 (STIMS bootstrap: *p* = 0.01) and Ara+2 (STIMS bootstrap: *p* = 0.035), although it is unclear whether the patterns in the nonmutator populations reflect relaxed or purifying selection (Grant, Maddamsetti, et al. 2021; Maddamsetti and Grant 2022).

**Figure 3.**
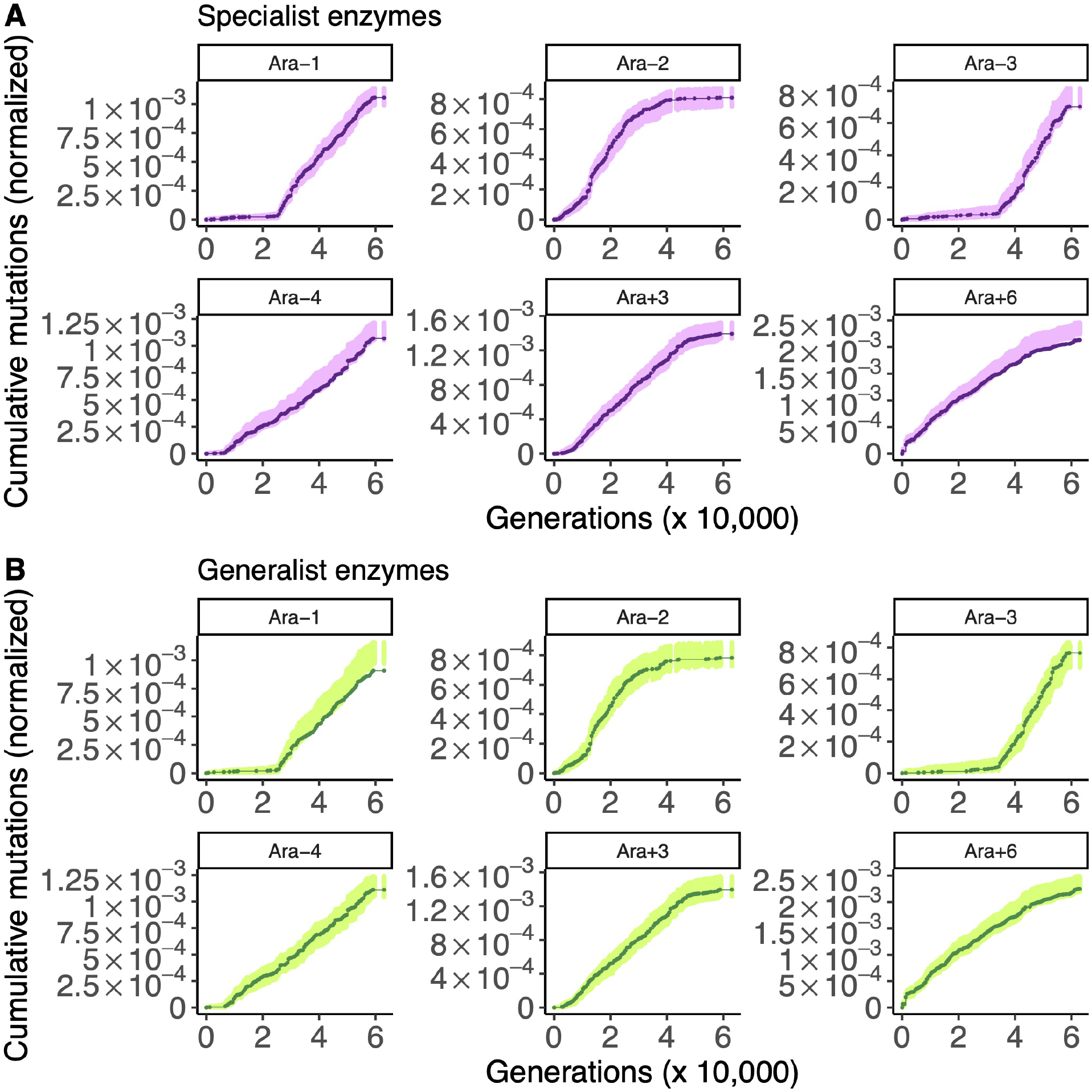
The results of running STIMS on specialist and generalist metabolic enzymes (hypermutator populations only). Each panel shows the cumulative number of mutations in the gene set of interest (solid line) in the six hypermutator LTEE populations. For comparison, random sets of genes (with the same cardinality as the gene set of interest) were sampled 1,000 times, and the cumulative number of mutations in those random gene sets, normalized by gene length, were calculated. The middle 95% of this null distribution is shown as shaded points. When a solid line falls below the shaded region, then the gene set of interest show a significant signal of purifying selection. A) The results of running STIMS on specialist enzymes B) The results of running STIMS on generalist enzymes.

Ara-1 (STIMS bootstrap: *p* = 0.001) is the only population in which I found evidence of purifying selection on generalist enzymes (Figure 3B, Supplementary Figure S2B). One nonmutator population, Ara+5 (Supplementary Figure S2B), shows evidence of positive selection on generalist enzymes (STIMS bootstrap: *p* = 0.0277). Altogether, these results somewhat support the hypothesis that specialist enzymes evolve under stronger purifying selection than generalist enzymes, with Ara-1 serving as a notable exception.

### The Jenga hypothesis can explain idiosyncratic variation in the strength of purifying selection on metabolism across replicate populations

What may account for this idiosyncratic variation? One possibility is that the strength of purifying selection on metabolic enzymes depends on *how* each population has lost metabolic function over time (Cooper and Lenski 2000; Leiby and Marx 2014). Ara+6 in particular has lost the ability to grow on many sugars that the ancestral LTEE clone can use (Leiby and Marx 2014). Those losses of function might have caused stronger purifying selection on the metabolic pathways that have remained intact. One way to conceptualize this hypothesis is as follows: at first, many metabolic pathways may evolve nearly neutrally, such that many loss-of-function mutations accumulate, especially in the hypermutator LTEE populations. Eventually, the accumulated effects of these losses of function reach a critical point, after which further gene losses become deleterious. Idiosyncratic differences in the strength of purifying selection on metabolic enzymes across populations then reflects idiosyncratic changes in the pathways that metabolic flux can follow in each population.

I call this conceptual model the Jenga hypothesis, after a children’s game in which wooden blocks are progressively removed from a tower until it falls. The Jenga hypothesis generalizes to any phenotype that is mechanistically caused by a molecular network with many redundancies (Albergante, et al. 2014). Such a network may have many nodes or edges that can be lost without any loss of function. At some point however, the network has lost most of its redundancies, causing strong purifying selection on the particular nodes that if lost would compromise network function. The specific nodes that end up evolving under strong purifying selection strongly depends on the particular history of nodes that were previously lost in the network, causing idiosyncratic purifying selection (Figure 4A).

**Figure 4.**
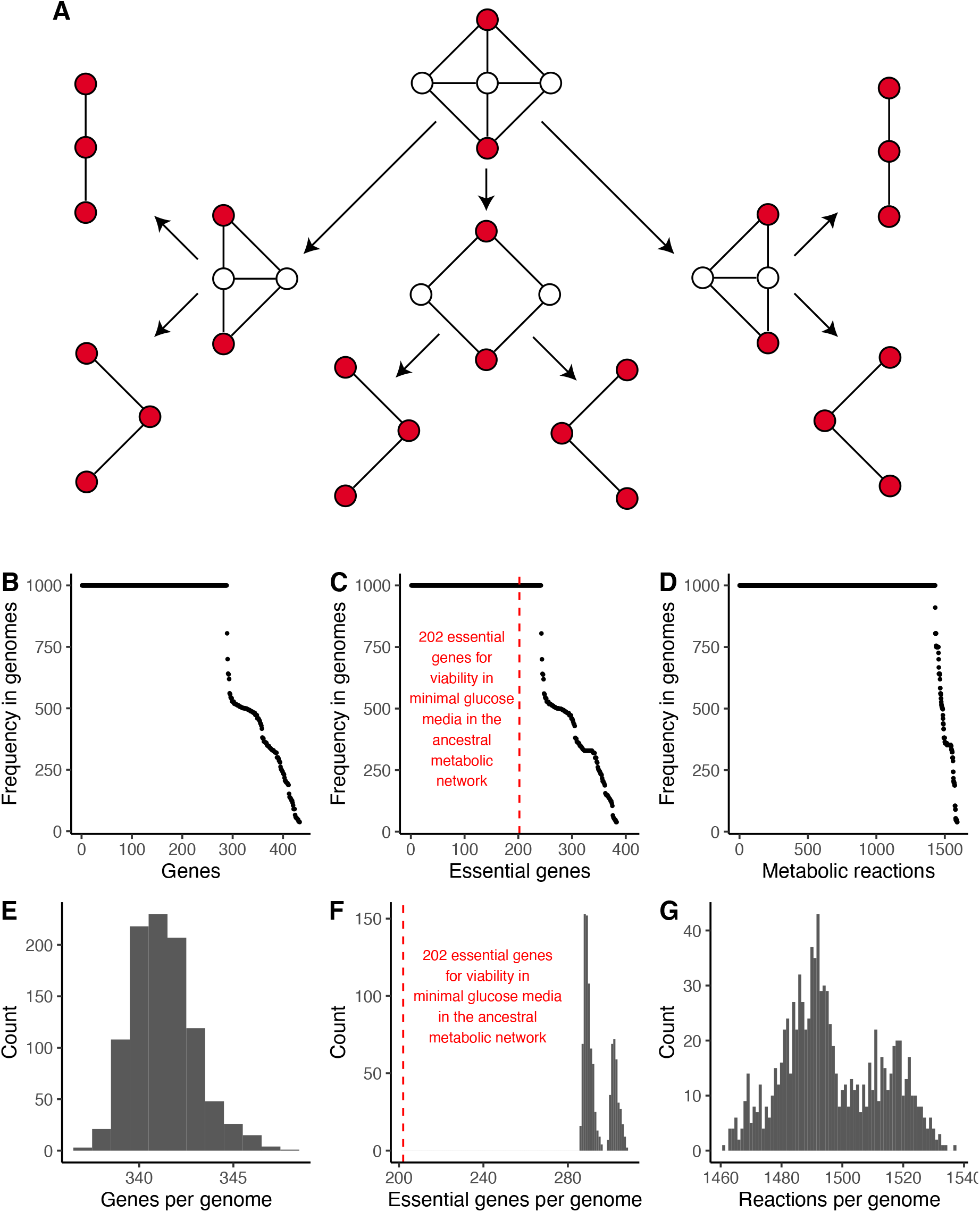
A computational model of network evolution under strong purifying selection demonstrates the Jenga hypothesis. A) The Jenga hypothesis explains how idiosyncratic purifying selection can arise in a network encoding an important phenotype. Suppose a given network is evolving under purifying selection, such that connections between red nodes must be maintained. Redundant intermediate nodes can be lost without affecting the network phenotype. Once all redundant nodes are lost, the remaining nodes evolve under strong purifying selection to maintain network function. The Jenga Hypothesis makes four predictions: 1) variation in minimal genome content 2) variation in minimal genome size 3) evolved networks are more fragile (more essential genes) 4) idiosyncratic variation in which particular genes are essential. All four predictions are supported by a model of metabolic network evolution in which 1,000 minimal genomes were evolved *in silico* under strong purifying selection for growth in minimal glucose media (Methods). B) Minimal genomes vary in gene content. C) Minimal genomes vary in essential gene content. Evolved genomes are more fragile (more essential genes) than the ancestral genome, indicated by the dashed red line D) Minimal genomes vary in metabolic reaction content. E) Minimal genomes have variable numbers of genes. F) Evolved genomes are more fragile (have more essential genes) than the ancestral genome, as indicated by the dashed red line, and show a bimodal distribution of essential genes, indicating the evolution of multiple network topologies under purifying selection. See Supplementary Figure 3 for details. G) The size distribution of the reaction networks encoded by the minimal genomes is multimodal, again indicating the evolution of multiple network topologies under purifying selection. See Supplementary Figure 3 for details.

The Jenga hypothesis makes four specific and testable predictions involving the reductive evolution of genomes encoding network phenotypes under purifying selection. First, replicate minimal genomes that have evolved under strong purifying selection should show variation in minimal genome content (i.e. the specific genes found in each genome). Second, replicate minimal genomes should have variable numbers of genes. Third, replicate minimal genomes should encode more fragile networks; in other words, they should have higher proportions of essential genes compared to their ancestor. Finally, replicate minimal genomes should show idiosyncratic variation in *which* particular genes are essential for network function. To test these predictions, I played 1,000 rounds of Genome Jenga, a zero-player game (like Conway’s game of Life) in which genes are randomly selected from a genome, and are removed if doing so has a negligible effect on fitness. For comparison to the LTEE, I played Genome Jenga in simulated minimal glucose media, and used the large effective population size (3.3 ×10^7^) of the LTEE (Izutsu, et al. 2021) to set the selective threshold (Methods).

All four predictions of the Jenga hypothesis are supported by these simulations (Figure 4). While there are core sets of genes (Figure 4B), essential genes (Figure 4C), and metabolic reactions (Figure 4D) found in all 1,000 minimal genomes, there are hundreds of genes (Figure 4B), essential genes (Figure 4C), and metabolic reactions (Figure 4D) which are found in some, but not all of the minimal genomes. The number of genes found in each minimal genome also varies (Figure 4E). All of the evolved metabolic networks encoded by the minimal genomes contain more essential genes—in absolute numbers— than the ancestral metabolic network (Figure 4C and Figure 4F). Strikingly, the distribution of the number of essential genes in the 1,000 minimal genomes is bimodal (Figure 4F), and the distribution of the number of metabolic reactions per genome is multimodal (Figure 4G). This indicates that several of the minimal genomes have converged to alternative network topologies with alternative sets of essential genes, which is further supported by how the minimal genomes cluster by essential gene content and metabolic reaction context (Supplementary Figure S3).

### Genes under purifying selection in the minimal genomes generated by Genome Jenga evolve under strong purifying selection across LTEE hypermutator populations

I then asked whether the gene context of the minimal genomes evolved through Genome Jenga could predict the metabolic pathways evolving under purifying selection in the LTEE. First, I examined the 288 core genes found in all of the 1,000 minimal genomes. 5 out of the 6 LTEE hypermutator populations show evidence of purifying selection on the core metabolic genes identified by Genome Jenga (Figure 5A): Ara-1 (STIMS bootstrap: *p* < 0.001), Ara-2 (STIMS bootstrap: *p* = 0.034), Ara-4 (STIMS bootstrap: *p* = 0.017), Ara+3 (STIMS bootstrap: *p* = 0.008), and Ara+6 (STIMS bootstrap: *p* < 0.001). Ara+4 is the sole nonmutator population (Supplementary Figure 4A) that shows a significant depletion of observed mutations in these core metabolic genes (STIMS bootstrap: *p* = 0.001). I also examined the 242 genes which are essential for viability on glucose in the 1,000 minimal genomes (Figure 5B). The same five hypermutator populations show evidence of purifying selection on these genes: Ara-1 (STIMS bootstrap: *p* < 0.001), Ara-2 (STIMS bootstrap: *p* = 0.023), Ara-4 (STIMS bootstrap: *p* = 0.053), Ara+3 (STIMS bootstrap: *p* = 0.003), and Ara+6 (STIMS bootstrap: *p* < 0.001). Two nonmutator populations (Supplementary Figure 4B) show a significant depletion of observed mutations in these core essential genes: Ara+1 (STIMS bootstrap: *p* = 0.017) and Ara+4 (STIMS bootstrap *p* = 0.005).

**Figure 5.**
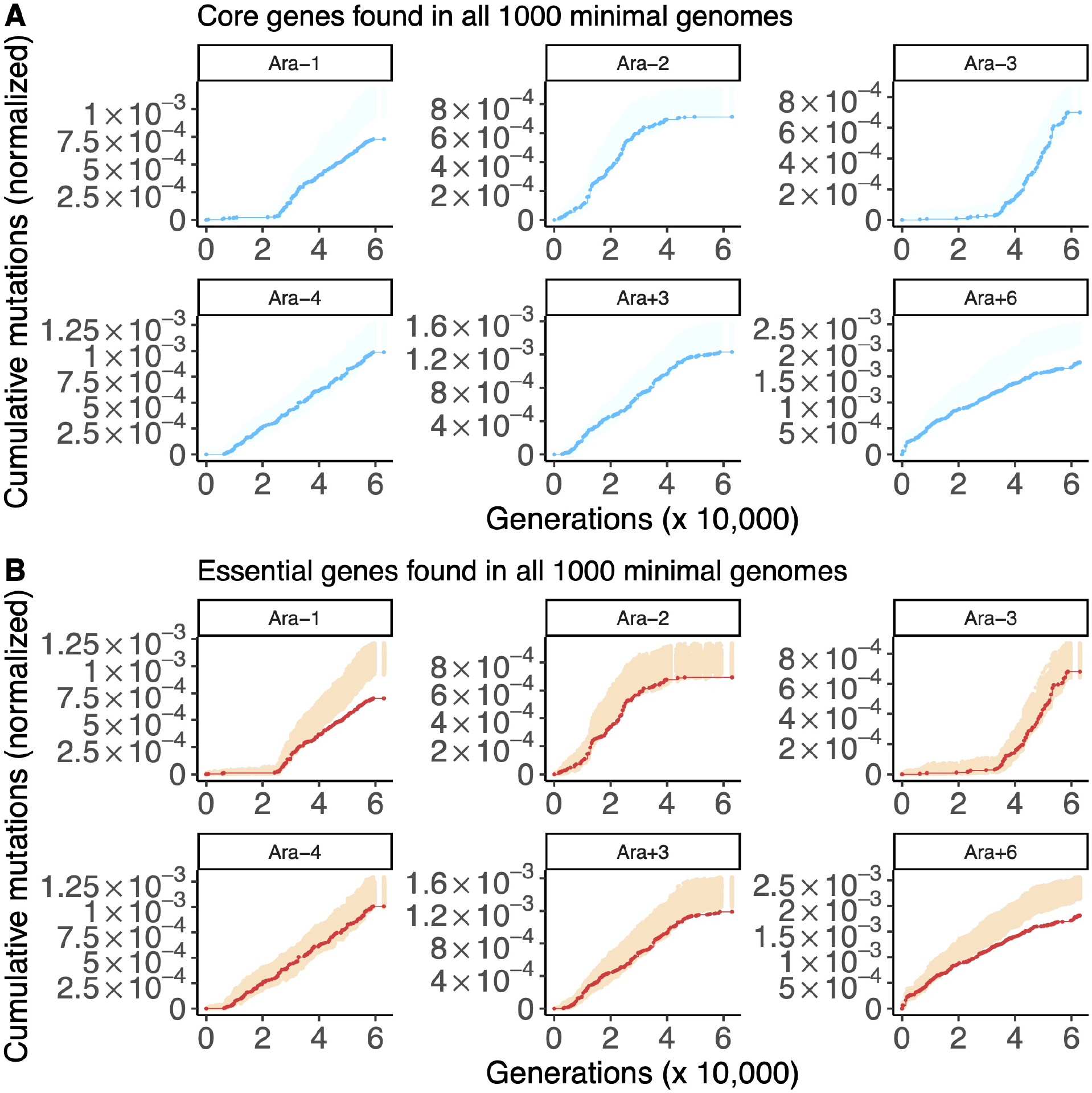
The results of running STIMS on core and essential genes in all 1,000 minimal genomes (hypermutator populations only). Each panel shows the cumulative number of mutations in the gene set of interest (solid line) in the six hypermutator LTEE populations. For comparison, random sets of genes (with the same cardinality as the gene set of interest) were sampled 1,000 times, and the cumulative number of mutations in those random gene sets, normalized by gene length, were calculated. The middle 95% of this null distribution is shown as shaded points. When a solid line falls below the shaded region, then the gene set of interest show a significant signal of purifying selection. A) The results of running STIMS on the core genes found in all 1,000 minimal genomes enzymes B) The results of running STIMS on the genes essential for viability on glucose in all 1,000 minimal genomes.

### Genes essential for viability on glucose or citrate in the ancestral REL606 metabolic model show strong purifying selection across LTEE hypermutator populations

Ara-3 is the only hypermutator population that does not show purifying selection on either the core or essential genes of the minimal genomes. Since Ara-3 evolved the ability to use citrate as a carbon source under aerobic conditions, a trait that none of the other LTEE populations have evolved (Blount, et al. 2008; Blount, et al. 2012), I hypothesized that Ara-3 would show purifying selection on genes that are predicted to be essential for viability on citrate by flux balance analysis (Ebrahim, et al. 2013). So, I asked whether the genes that are predicted to be essential for growth on glucose or citrate in the ancestral REL606 metabolic model showed evidence of purifying selection in the LTEE hypermutator populations (Figure 6).

**Figure 6.**
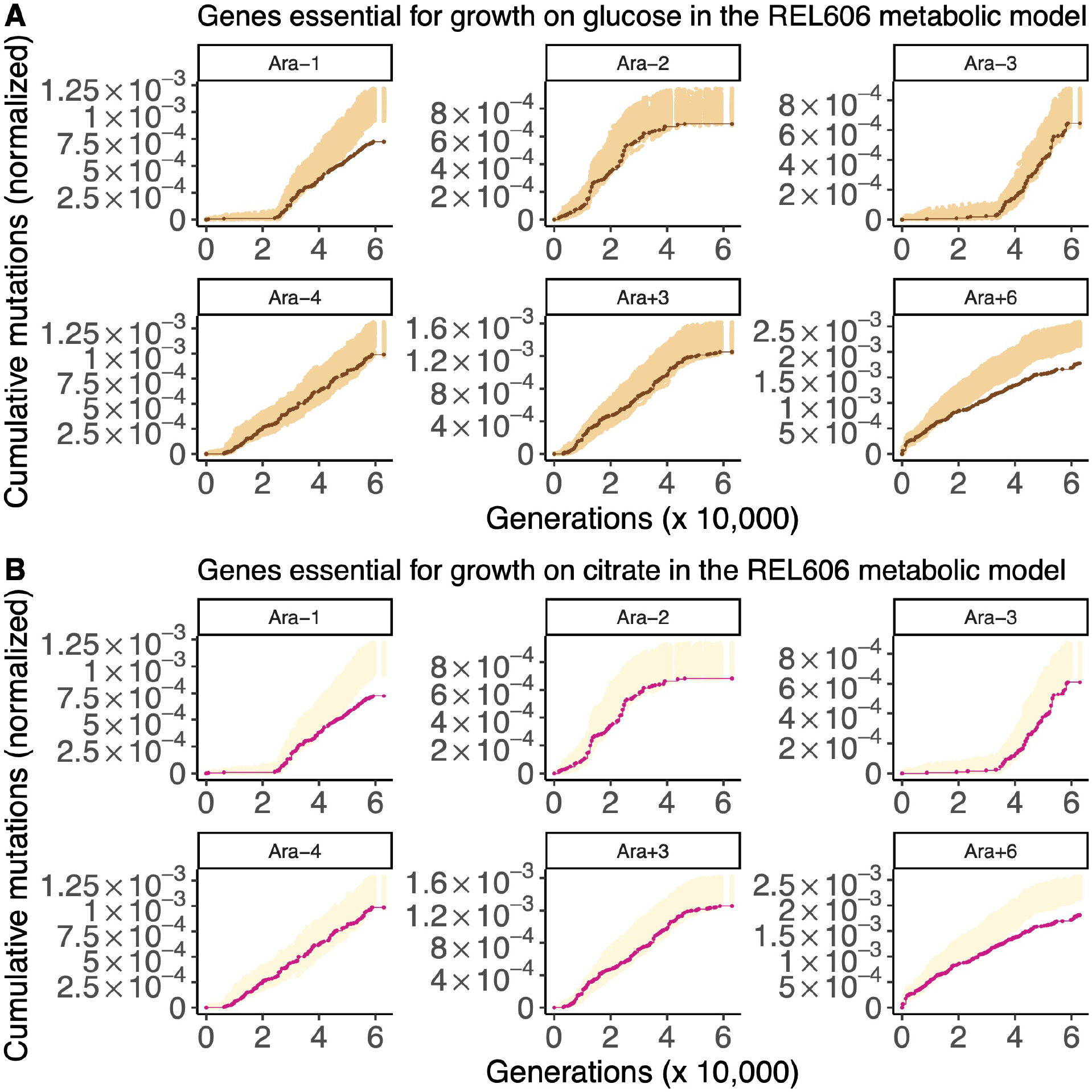
The results of running STIMS on essential genes for growth on either glucose or citrate in the ancestral REL606 metabolic model (hypermutator populations only). Each panel shows the cumulative number of mutations in the gene set of interest (solid line) in the six hypermutator LTEE populations. For comparison, random sets of genes (with the same cardinality as the gene set of interest) were sampled 1000 times, and the cumulative number of mutations in those random gene sets, normalized by gene length, were calculated. The middle 95% of this null distribution is shown as shaded points. When a solid line falls below the shaded region, then the gene set of interest show a significant signal of purifying selection. A) The results of running STIMS on the genes essential for viability on glucose in the REL606 metabolic model. B) The results of running STIMS on the genes essential for viability on citrate in the REL606 metabolic model.

For genes predicted to be essential on glucose (Figure 6A), all 6 hypermutator populations show signs of purifying selection (Ara-1:*p* < 0.001; Ara-2: *p* = 0.035; Ara-3: 0.051; Ara-4: *p* = 0.053; Ara+3: *p* = 0.051; Ara+6: *p* < 0.001). Ara+1 and Ara+4 are again the two nonmutator populations depleted of mutations, as shown in Supplementary Figure 5A (Ara+1 STIMS bootstrap: *p* = 0.047; Ara+4: *p* < 0.001).

For genes predicted to be essential on citrate (Figure 6B), all 6 hypermutator populations show signs of purifying selection (Ara-1: *p* < 0.001; Ara-2:*p* = 0.025; Ara-3: 0.0139; Ara-4:*p* = 0.050; Ara+3: *p* = 0.057; Ara+6: *p* < 0.001). Ara+1 and Ara+4 are again the two nonmutator populations depleted of mutations, as shown Supplementary Figure 5B (Ara+1: *p* = 0.040; Ara+4: *p* < 0.001).

The sets of genes are that essential for growth on glucose or citrate in the ancestral metabolic model, and the sets of genes that are essential or found in all of the 1,000 minimal genomes show considerable overlap: of the 203 genes predicted to be essential for growth on citrate, only 4 are predicted to be non-essential for growth on glucose (Figure 7). In this light, it is especially notable that the Cit^+^ Ara–3 hypermutator population shows a much stronger signal of purifying selection on genes that are predicted to be essential for growth on citrate in the REL606 genome-scale metabolic model.

**Figure 7.**
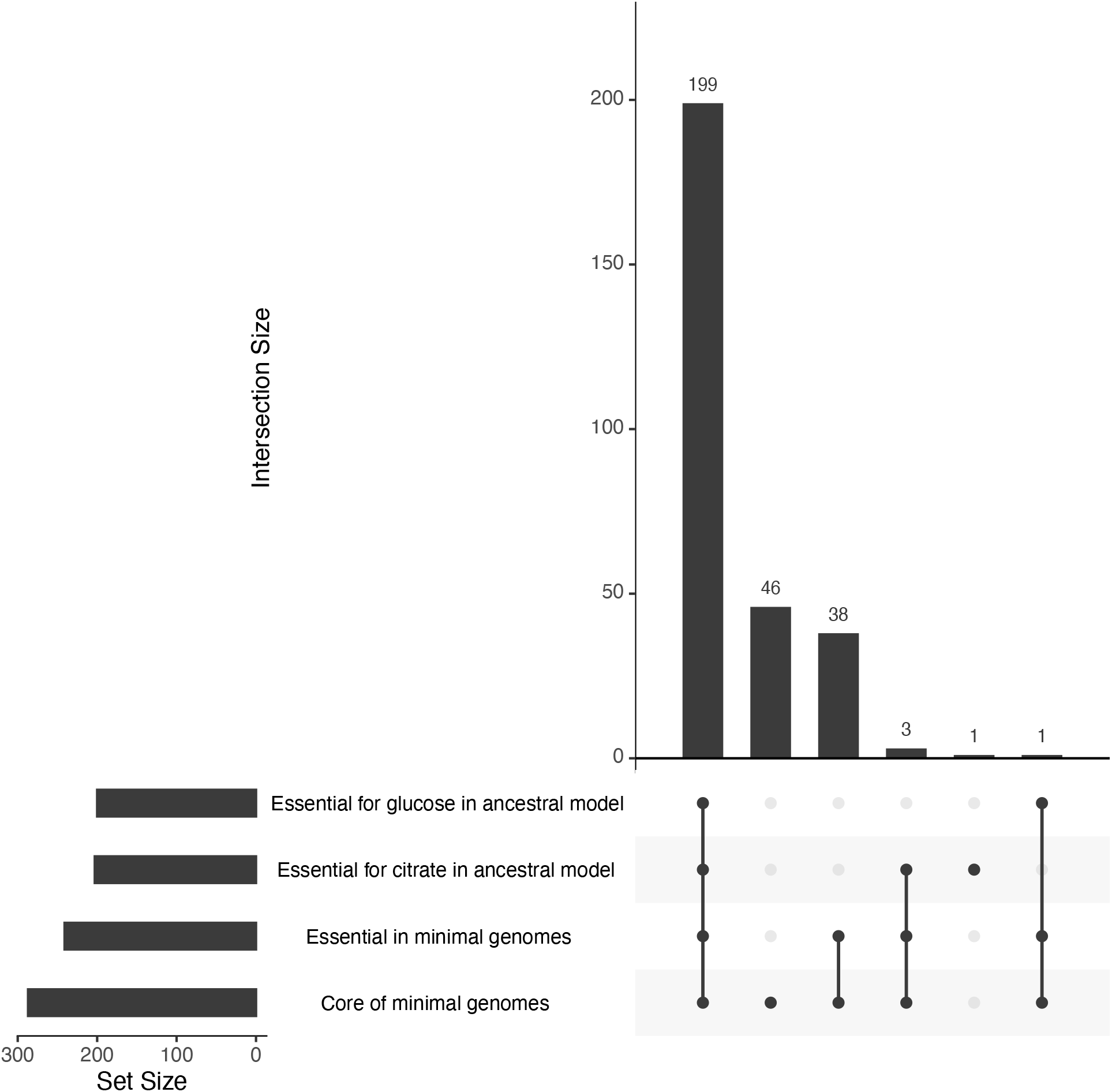
Intersections among the sets of metabolic enzymes defined by playing Genome Jenga. The intersections between four sets were examined: core enzymes among the 1,000 minimal genomes, essential enzymes among the 1,000 minimal genomes, essential enzymes for growth on glucose in the ancestral REL606 metabolic model, and essential enzymes for growth on citrate in the ancestral REL606 metabolic model. The cardinality of each of these sets is shown in the horizontal bar graph on the left. Every non-empty subset of the four sets is visualized as a set of dots connected by lines. The cardinality for each non-empty subset (set intersections) is shown in the vertical bar graph.

## Discussion

I tested four sets of metabolic genes for purifying selection in the LTEE: those encoding reactions in central metabolism; those catalyzing “superessential” reactions, and those that encode specialist and generalist enzymes in the *E. coli* metabolic reaction network. Evidence of purifying selection was found only in specific populations. This finding indicates that the strength of purifying selection on metabolism in each population depends on its particular evolutionary history during the LTEE. In this regard, it is important to note that two hypermutator populations may be considered as outliers for the purpose of this paper. First, Ara–2 lost its hypermutator phenotype following the fixation of a reversion mutation at 42,250 generations (Maddamsetti and Grant 2020). It may be challenging to detect a cumulative signal of purifying selection in later generations of this population owing to the lower mutation rate (Maddamsetti and Grant 2022). Second, Ara–3 evolved the ability to use citrate as a carbon source at ~31,000 generations (Blount, et al. 2008; Blount, et al. 2012). Its phenotypic evolution is now on a different trajectory from the other populations (Blount, et al. 2018; Blount, et al. 2020; Grant, Abdel Magid, et al. 2021).

Overall, Ara+6 shows the strongest signal of purifying selection on metabolic enzymes. This finding is consistent with previous results that show strong purifying selection on aerobic and anaerobic-specific genes in Ara+6 (Grant, Maddamsetti, et al. 2021) and strong purifying selection on essential genes in Ara+6 (Maddamsetti and Grant 2022). Ara+6 has accumulated the most mutations over the course of the LTEE, which may explain its especially strong signal of purifying selection.

Nonetheless, mutation accumulation does not seem to be sufficient to explain the varying signal of purifying selection across populations. If mutation accumulation were a dominant factor across populations, then the populations with the most mutations should show the strongest signal of purifying selection on metabolic enzymes. However, like Ara+6, Ara+3 has also accumulated many mutations (Tenaillon, et al. 2016; Couce, et al. 2017; Maddamsetti and Grant 2020), and yet it shows much weaker evidence of purifying selection on metabolic enzymes. Moreover, Ara-1 shows by far the strongest signal of purifying selection on superessential genes, even though it evolved a hypermutator phenotype much later than Ara+3 and Ara+6 (Barrick and Lenski 2009; Maddamsetti, et al. 2015; Tenaillon, et al. 2016; Good, et al. 2017; Maddamsetti and Grant 2020).

To explain these idiosyncratic differences in purifying selection on metabolic enzymes, I proposed the Jenga hypothesis. The Jenga hypothesis is closely related to the hypothesis that interconnections between aerobic and anaerobic metabolism prevent the loss of anaerobic function despite evolving under relaxed selection in LTEE (Grant, Maddamsetti, et al. 2021). The main difference between the buttressing pleiotropy model proposed by Grant et al. (2021) and the model considered here is that I propose that purifying selection on metabolic enzymes may only become evident after passing a critical threshold of network evolution.

I tested the plausibility of the Jenga hypothesis by asking whether distinct reaction networks would evolve from an ancestral genome-scale metabolic model of the REL606 genome. By simulating genome evolution under strong purifying selection for growth on minimal glucose media, I found clear evidence for multiple predictions that follow from the Jenga hypothesis. While the minimal genomes share a core set of genes and reactions, they broadly vary in terms of gene content, essential genes, metabolic reactions. In particular, more genes became essential in all 1,000 minimal genomes, and each minimal genome has an idiosyncratic set of genes that is required for growth in simulated minimal glucose media.

In addition, the core genes found across the minimal genomes show clear and consistent signatures of purifying selection across the LTEE populations. This finding suggests that the subset of metabolic enzymes that are absolutely required for growth on glucose (or on citrate in the case of Ara-3) is evolving under purifying selection across the LTEE. Together with my initial findings on core metabolism, superessential genes, and specialist and generalist enzymes, my results imply that the large number of metabolic redundancies present in the *E. coli* genome during growth on a small number of carbon sources, especially given the static and unchanging nature of the laboratory environment, can result in divergent network evolution during an ongoing process of genome streamlining. Future research could study the implications of the Jenga hypothesis using evolution experiments with organisms with minimal genomes, including endosymbionts and engineered organisms (Gil, et al. 2002; Hutchison, et al. 2016). Specifically, organisms with minimal genomes should show strong purifying selection against losses of metabolic function, while adding redundancies should weaken purifying selection.

The Jenga hypothesis makes a further, specific prediction with regard to the LTEE. The function of many metabolic genes, or their regulation, may already be knocked out in many populations given the extensive losses of metabolic function that have been observed (Leiby and Marx 2014). In most cases, the genetic causes of these losses of function are unknown. If idiosyncratic purifying selection is caused by the progressive loss of metabolic function, then one should be able to predict which enzymes have come under the strongest purifying selection in the network, based on which enzymes have the greatest gains in metabolic flux over adaptive evolution, or the least variability in (or equivalently, the most constraint on) metabolic flux. Testing this prediction will require metabolomics experiments to describe how metabolic flux has evolved in the LTEE, and targeted knockout experiments to see whether evolved bottlenecks in each population’s metabolic network can account for idiosyncratic purifying selection on metabolic enzymes in the LTEE.

## Materials and Methods

### Data sources

Pre-processed LTEE metagenomic data were downloaded from: https://github.com/benjaminhgood/LTEE-metagenomic. Genes in the BiGG *E. coli* core metabolic model were downloaded from: http://bigg.ucsd.edu/models/e_coli_core. *E. coli* genes encoding superessential metabolic reactions were taken from supplementary table S6 of Barve et al. (2014). *E. coli* genes encoding specialist and generalist metabolic enzymes were taken from supplementary table S1 of Nam et al. (2012). These tables were then merged with NCBI Genbank gene annotation for the ancestral LTEE strain, *Escherichia coli* B str. REL606. The UpSet visualization for the overlap between the four sets of metabolic enzymes was produced using the UpSetR R Package (Conway, et al. 2017).

### Data analysis with STIMS

STIMS is fully described in Maddamsetti and Grant (2022). Briefly, STIMS counts the cumulative number of mutations occurring over time in a gene set of interest, and compares that number to a null distribution that is constructed by subsampling random gene sets of equivalent size. The number of observed mutations in a gene set is normalized by the total length of that gene set in nucleotides. Bootstrapped *p*-values were calculated separately for each population. *p*-values for one-sided tests for purifying selection were calculated as the lower-tail probability, assuming the null distribution, of sampling a normalized cumulative number of mutations that is less than the normalized cumulative number of mutations in the gene set of interest.

In the visualizations shown in the figures, the top 2.5% and bottom 2.5% of points in the null distribution are omitted, such that each panel can be interpreted as a two-sided randomization test with a false-positive (type I error) probability α = 0.05.

### Genome Jenga

In Genome Jenga, genes are randomly selected from a genome, and removed if doing so has a negligible effect on fitness. The game is continued until no further genes can be removed from the genome. I used a genome-scale metabolic model of the ancestral LTEE REL606 strain (King, et al. 2015), and I used the same algorithm for minimal genome construction reported by (Pál, et al. 2006). I generated 1,000 minimal genomes from the metabolic model of REL606, evaluating changes in fitness as the change in the predicted biomass objective optimized by flux balance analysis after the removal of a given gene (Pál, et al. 2006; Hosseini and Wagner 2018). I modeled the growth conditions of the LTEE by using glucose as the sole limiting carbon source, and adding an excess of thiamine to model the Davis-Mingioli medium used in the LTEE (Lenski, et al. 1991). I modeled strong purifying selection by setting the fitness threshold *s* for gene removal as *1/N_e_,* where *N_e_* = 3.3 × 10^7^, which is the effective population size of the LTEE (Izutsu, et al. 2021). This models nearly-neutral evolution in which mutations that disrupt a gene’s function can hitchhike to fixation if they either confer a beneficial effect, or confer a sufficiently small fitness defect such that |*N_e_ s*| < 1, where *s* is the selective effect of the gene disruption (Pál, et al. 2006). My design deviates from that in Pál et al. (2006) in that I model reductive genome evolution in minimal glucose media rather than rich media, and I used a highly stringent selective threshold to model the large population size of the LTEE in contrast to the small effective population sizes of intracellular endosymbionts (*N_e_* ~ 10^2^–10^3^) modeled by Pál et al. (2006). I used COBRAPy software to conduct flux balance analyses, including calculations of which genes in the metabolic model were essential for growth (Ebrahim, et al. 2013).

## Data Availability Statement

The data and analysis codes underlying this article are available on the Zenodo Digital Repository (DOI pending publication). Analysis codes are also available at: https://github.com/rohanmaddamsetti/LTEE-network-analysis.

## Acknowledgements

I thank Richard Lenski, Jeffrey Barrick, and Lingchong You for guidance and advice, Nkrumah Grant for valuable discussions and comments, and Zachary Blount for detailed comments and feedback on an earlier version of the manuscript. The LTEE is supported by a grant from the National Science Foundation (currently DEB-1951307) to Richard Lenski and Jeffrey Barrick.

## SUPPLEMENTARY INFORMATION

**Supplementary File 1: Mapping of REL606 genes into the four gene sets described in Figure 1**.

**Supplementary File 2: Mapping of REL606 genes into the four gene sets described in Figure 7**.

**Supplementary Figure 1.**
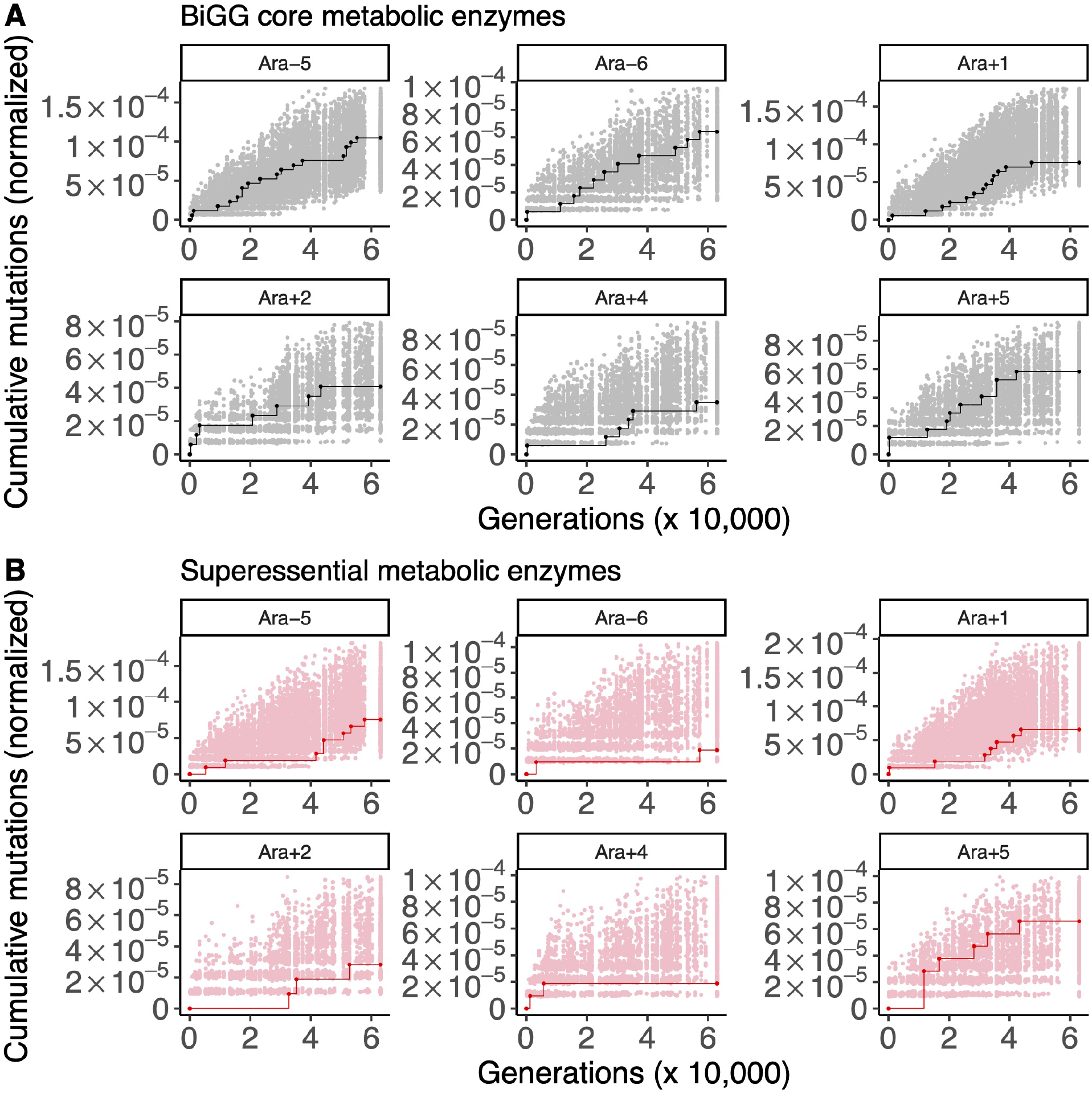
The results of running STIMS on BiGG core enzymes and superessential enzymes (nonmutator populations only). Each panel shows the cumulative number of mutations in the gene set of interest (solid line) in the six nonmutator LTEE populations. For comparison, random sets of genes (with the same cardinality as the gene set of interest) were sampled 1,000 times, and the cumulative number of mutations in those random gene sets, normalized by gene length, were calculated. The middle 95% of this null distribution is shown as shaded points. When a solid line falls above the shaded region, then the gene set of interest show a significant signal of positive selection. A) The results of running STIMS on enzymes in the BiGG *E. coli* core metabolism model. B) The results of running STIMS on enzymes catalyzing superessential metabolic reactions.

**Supplementary Figure 2.**
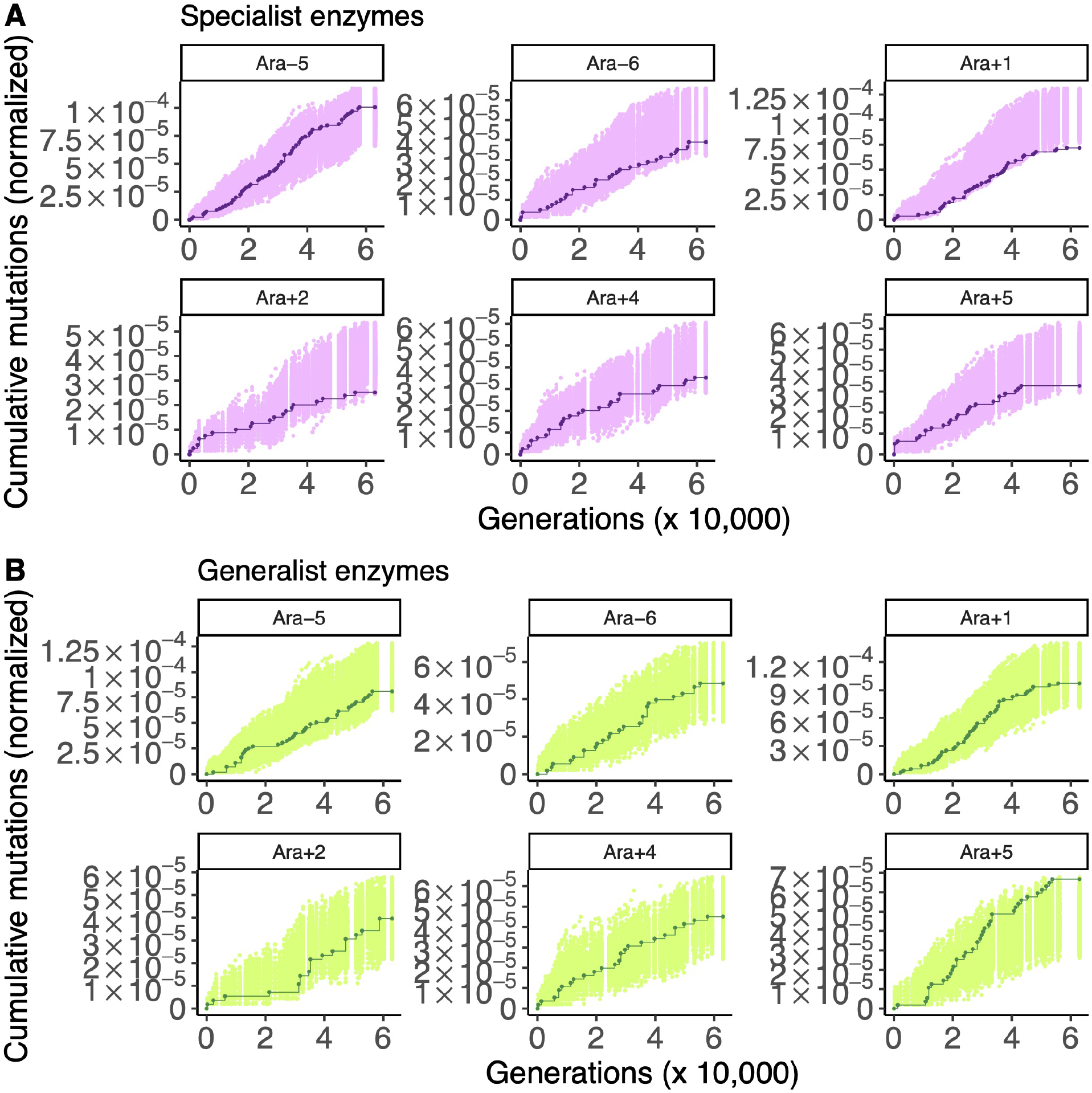
The results of running STIMS on specialist enzymes and generalist enzymes (nonmutator populations only). Each panel shows the cumulative number of mutations in the gene set of interest (solid line) in the six nonmutator LTEE populations. For comparison, random sets of genes (with the same cardinality as the gene set of interest) were sampled 1000 times, and the cumulative number of mutations in those random gene sets, normalized by gene length, were calculated. The middle 95% of this null distribution is shown as shaded points. When a solid line falls above the shaded region, then the gene set of interest show a significant signal of positive selection. A) The results of running STIMS on specialist enzymes. B) The results of running STIMS on generalist enzymes.

**Supplementary Figure 3.**
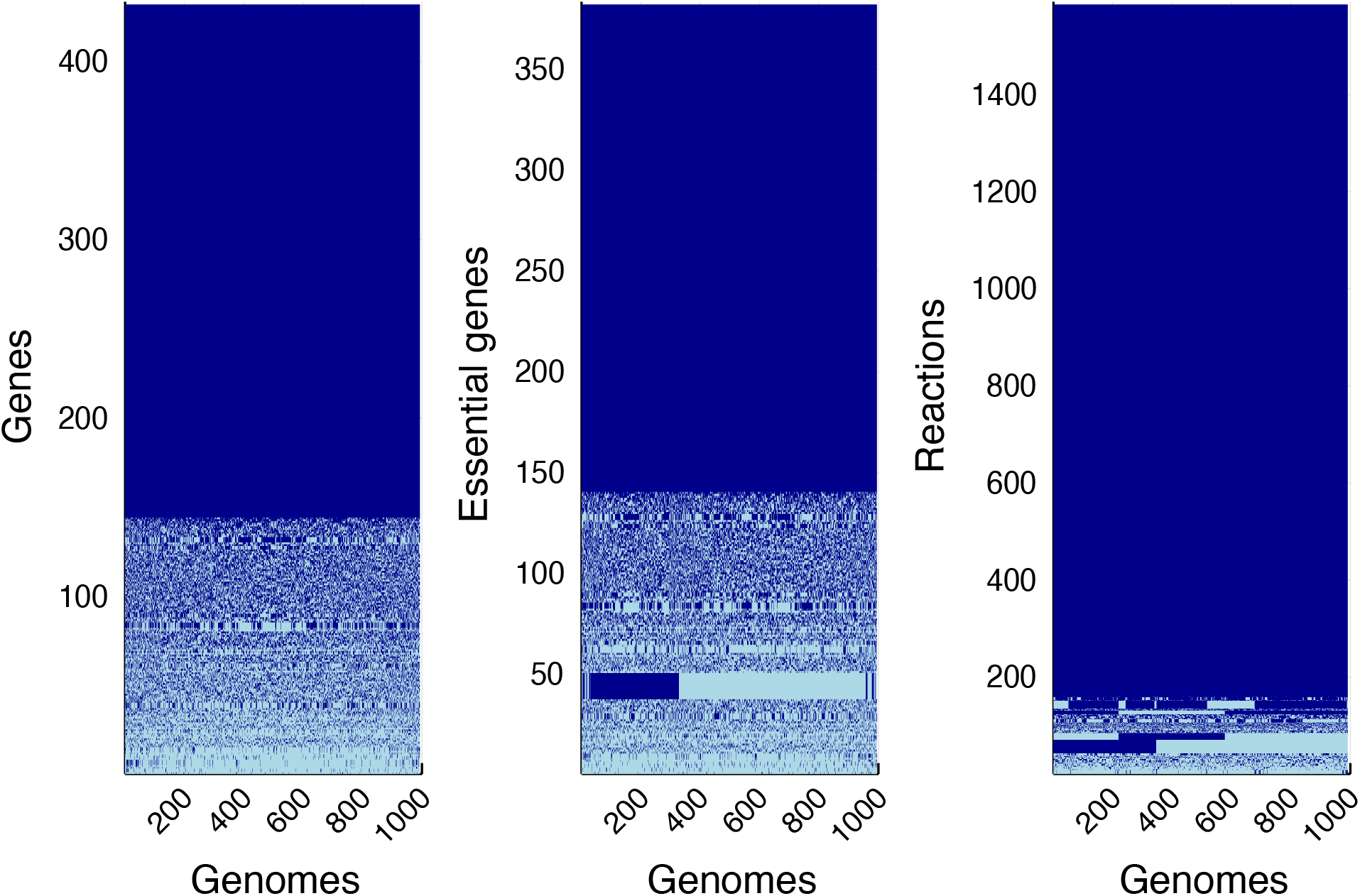
The block structure of genome content across 1,000 minimal genomes shows that strong purifying selection results in many possible minimal metabolic networks, as predicted by the Jenga Hypothesis. The genes and metabolic reactions present in each genome are shown in dark blue; light blue indicates absence. Core genes and reactions found in all 1,000 genomes account for the dark blue blocks seen in each panel. Left panel: matrix of genes in each minimal genome. Middle panel: matrix of genes essential for viability on glucose in each minimal genome. Right panel: matrix of metabolic reactions encoded by each minimal genome.

**Supplementary Figure 4.**
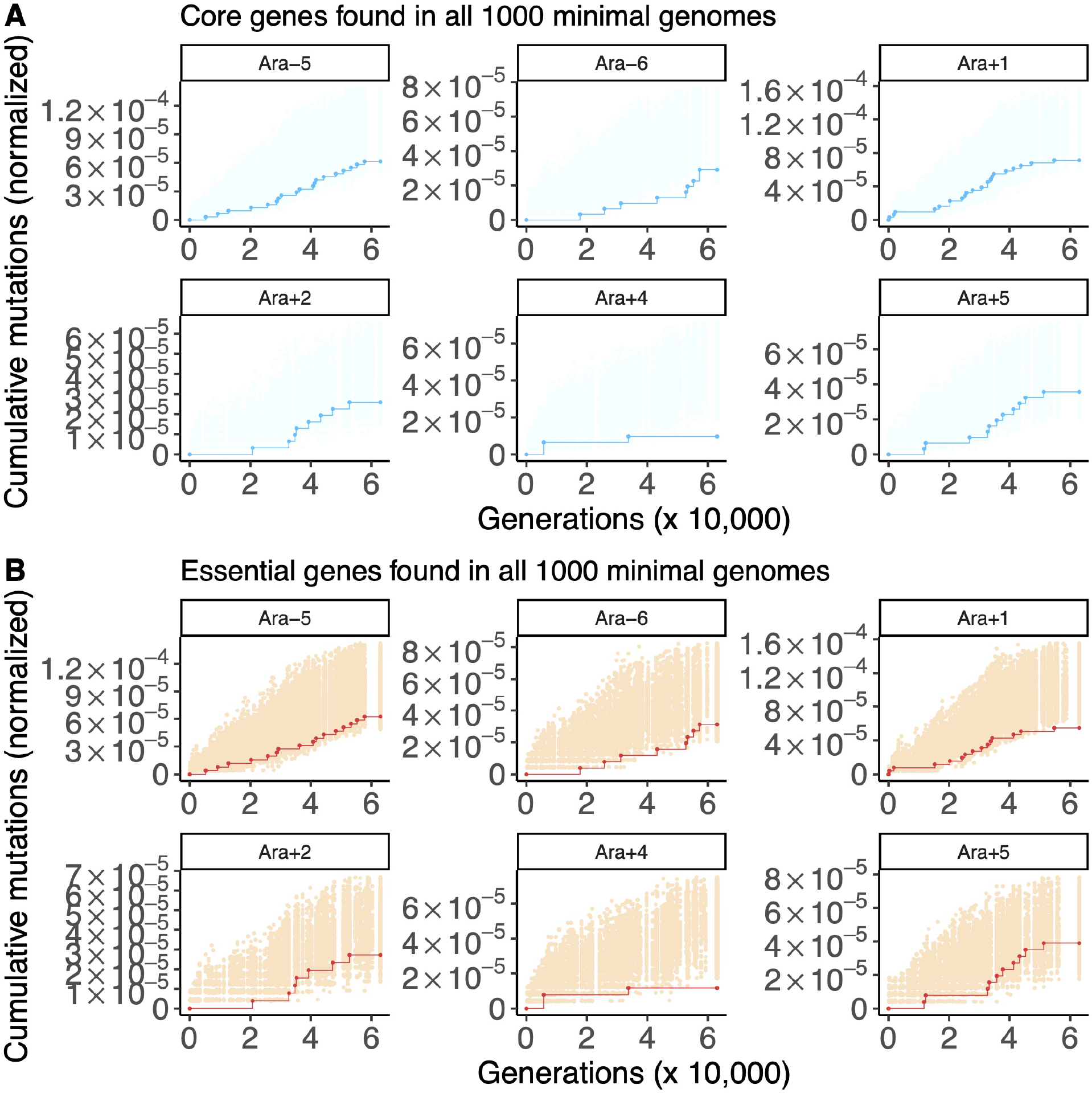
The results of running STIMS on core and essential genes in all 1,000 minimal genomes (nonmutator populations only). Each panel shows the cumulative number of mutations in the gene set of interest (solid line) in the six nonmutator LTEE populations. For comparison, random sets of genes (with the same cardinality as the gene set of interest) were sampled 1,000 times, and the cumulative number of mutations in those random gene sets, normalized by gene length, were calculated. The middle 95% of this null distribution is shown as shaded points. When a solid line falls below the shaded region, then the gene set of interest show a significant signal of purifying selection. A) The results of running STIMS on the core genes found in all 1,000 minimal genomes enzymes B) The results of running STIMS on the genes essential for viability on glucose in all 1000 minimal genomes.

**Supplementary Figure 5.**
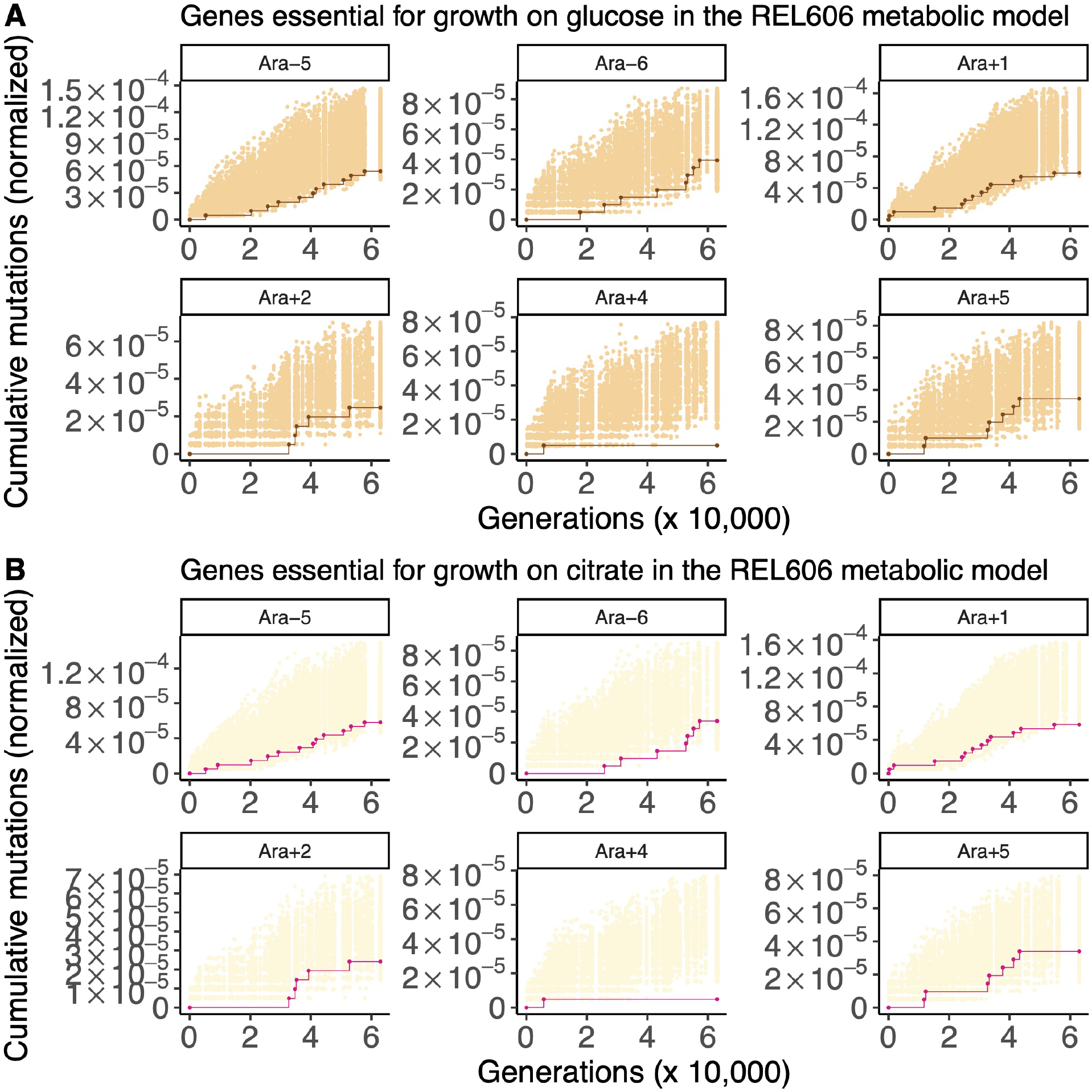
The results of running STIMS on essential genes for growth on either glucose or citrate in the ancestral REL606 metabolic model (nonmutator populations only). Each panel shows the cumulative number of mutations in the gene set of interest (solid line) in the six nonmutator LTEE populations. For comparison, random sets of genes (with the same cardinality as the gene set of interest) were sampled 1,000 times, and the cumulative number of mutations in those random gene sets, normalized by gene length, were calculated. The middle 95% of this null distribution is shown as shaded points. When a solid line falls below the shaded region, then the gene set of interest show a significant signal of purifying selection. A) The results of running STIMS on the genes essential for viability on glucose in the REL606 metabolic model. B) The results of running STIMS on the genes essential for viability on citrate in the REL606 metabolic model.

## Notes

### Competing Interest Statement

The authors have declared no competing interest.

